## Supplementary figures and images for "High throughput detection and genetic epidemiology of SARS-CoV-2 using COVIDSeq next generation sequencing"

### Supplementary Figure 1

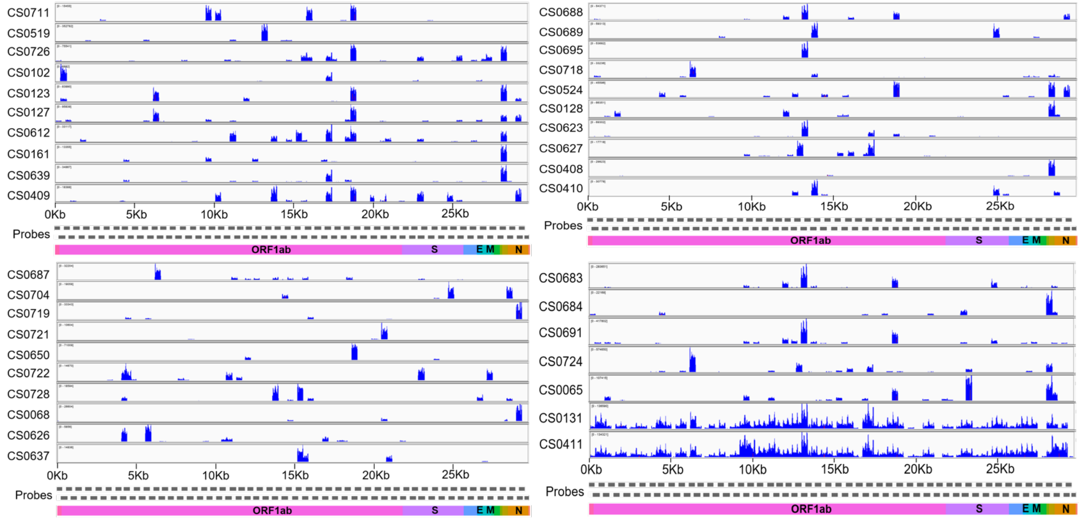
