## Supplementary Table 2 for "High throughput detection and genetic epidemiology of SARS-CoV-2 using COVIDSeq next generation sequencing"

| Task_name | duration (min:sec) | Duration (seconds) |
|---|---|---|
| LookForFastqFolderInCovid19SeqTask | 00:00.5 | 0.57 |
| LookForFastqFolderInValidationTask | 00:02.3 | 2.40 |
| SampleSheetValidationTask | 03:27.6 | 207.66 |
| RunQcTask | 02:40.7 | 160.79 |
| ResetFPGATask | 00:15.6 | 15.70 |
| FindDragenLicenseInstanceLocationTask | 00:05.0 | 5.01 |
| LookForFastqFolderInAnalysisTask | 00:05.0 | 5.01 |
| FindDragenLicenseTask | 00:03.9 | 3.98 |
| FirstTileOnlyTask | 00:03.9 | 3.98 |
| SingleLaneOnlyTask | 00:03.9 | 3.98 |
| FastModeTask | 00:03.9 | 3.98 |
| DragenFastqGenerationTask | 27:51.0 | 1671.07 |
| DragenKmerTask | 20:23.7 | 1223.77 |
| CopyTargetBedTask | 00:02.9 | 2.92 |
| FindSampleValidityTask | 05:56.9 | 356.94 |
| DragenMapAlignTask | 29:16.8 | 8956.87 |
| DragenVariantCallingTask | 26:15.1 | 8775.14 |
| ConsensusFastaTask | 09:37.0 | 577.01 |
| ReportGateKeeperTask | 00:01.3 | 1.30 |
| JsonTsvReportTask | 07:46.0 | 466.09 |
| StartReportEngineTask | 00:12.1 | 12.18 |
| RenderReportTask | 00:08.7 | 8.75 |
| StopReportEngineTask | 00:00.5 | 0.57 |
| DsdmErrorSummaryTask | 00:02.5 | 2.52 |
| Total Time (in seconds) | | 22468.19 |
| Total Time (in minutes) | | 374.47 |
| Total Time (in hours) | | 6.24 |
