## Supplementary Table 3 for "High throughput detection and genetic epidemiology of SARS-CoV-2 using COVIDSeq next generation sequencing"

| | | RT-PCR | | DRAGEN COVIDSeq Test Pipeline | | In - Depth Custom Pipeline (Merged) | | | | | | | | | | | | | | |
|---|---|---|---|---|---|---|---|---|---|---|---|---|---|---|---|---|---|---|---|---|
| Sample IDs Replicate 1 | Sample IDs Replicate 2 | RT (Pos/Neg) replicate1 | RT (Pos/Neg) replicate 2 | DRAGEN SARS-CoV-2 Detection Replicate 1 | DRAGEN SARS-CoV-2 Detection Replicate 2 | Total Reads_merged | Trimmed Reads q30-30bp_merged | Hisat2 Human % _merged | Hisat2 Covidall % _merged | Hisat2 Unmapped Covid Read#_merged | Hisat2 Unmapped Covid %_merged | BasePairescovered_merged | X coverage _merged | Genome coverage_merged | # bases with 0 coverage_merged | # bases with >0 coverage_merged | # bases with <10x coverage_merged | # bases with <100x coverage_merged | VARSCAN Variant Count_merged | N's_merged |
| CS0531_1 | CS0531_2 | Positive | Positive | SARS-CoV-2 Detected | SARS-CoV-2 Detected | 18352585 | 17194961 | 1.11 | 98.08 | 17003620 | 99.19 | 599305340 | 20041.64599 | 99.96321439 | 11 | 29892 | 56 | 88 | 13 | 57 |
| CS0530_1 | CS0530_2 | Positive | Positive | SARS-CoV-2 Detected | SARS-CoV-2 Detected | 14110540 | 13190264 | 4.28 | 94.65 | 12625755 | 98.89 | 443779374 | 14840.63051 | 99.86289001 | 41 | 29862 | 49 | 241 | 15 | 206 |
| CS0529_1 | CS0529_2 | Positive | Positive | SARS-CoV-2 Detected | SARS-CoV-2 Detected | 17064503 | 16003804 | 4.27 | 94.67 | 15320304 | 98.89 | 538524076 | 18009.03174 | 99.85954586 | 42 | 29861 | 60 | 259 | 14 | 234 |
| CS0533_1 | CS0533_2 | Positive | Positive | SARS-CoV-2 Detected | SARS-CoV-2 Detected | 16363129 | 15369240 | 1.1 | 97.96 | 15200834 | 99.04 | 535287906 | 17900.80948 | 99.78263051 | 65 | 29838 | 81 | 251 | 13 | 311 |
| CS0545_1 | CS0545_2 | Positive | Positive | SARS-CoV-2 Detected | SARS-CoV-2 Detected | 19052128 | 17733506 | 1.04 | 98.03 | 17548399 | 99.06 | 617693386 | 20656.56911 | 99.95987025 | 12 | 29891 | 58 | 78 | 12 | 67 |
| CS0532_1 | CS0532_2 | Positive | Positive | SARS-CoV-2 Detected | SARS-CoV-2 Detected | 19216801 | 17895898 | 7.34 | 91.17 | 16581942 | 98.4 | 579840423 | 19390.71073 | 99.96990269 | 9 | 29894 | 57 | 99 | 14 | 259 |
| CS0544_1 | CS0544_2 | Positive | Positive | SARS-CoV-2 Detected | SARS-CoV-2 Detected | 20077876 | 18870898 | 0.93 | 98.21 | 18694583 | 99.14 | 658874134 | 22033.71347 | 99.95987025 | 12 | 29891 | 56 | 198 | 10 | 317 |
| CS0543_1 | CS0543_2 | Positive | Positive | SARS-CoV-2 Detected | SARS-CoV-2 Detected | 19486012 | 18416819 | 4.68 | 94.27 | 17555450 | 98.89 | 617222854 | 20640.83383 | 99.85620172 | 43 | 29860 | 76 | 238 | 15 | 300 |
| CS0542_1 | CS0542_2 | Positive | Positive | SARS-CoV-2 Detected | SARS-CoV-2 Detected | 14463699 | 13604022 | 8.34 | 89.96 | 12469734 | 98.14 | 435003119 | 14547.13972 | 99.96655854 | 10 | 29893 | 50 | 115 | 15 | 332 |
| CS0541_1 | CS0541_2 | Positive | Positive | SARS-CoV-2 Detected | SARS-CoV-2 Detected | 20335078 | 18823699 | 2.99 | 95.95 | 18260096 | 98.91 | 641771847 | 21461.78801 | 99.96321439 | 11 | 29892 | 52 | 82 | 15 | 60 |
| CS0540_1 | CS0540_2 | Positive | Positive | SARS-CoV-2 Detected | SARS-CoV-2 Detected | 18167494 | 17053272 | 3.3 | 95.75 | 16490328 | 99.02 | 580458127 | 19411.36766 | 99.94649366 | 16 | 29887 | 55 | 133 | 12 | 262 |
| CS0539_1 | CS0539_2 | Positive | Positive | SARS-CoV-2 Detected | SARS-CoV-2 Detected | 17827645 | 16711870 | 1.43 | 97.77 | 16472549 | 99.19 | 580858888 | 19424.76969 | 99.85285757 | 44 | 29859 | 69 | 90 | 10 | 258 |
| CS0538_1 | CS0538_2 | Positive | Positive | SARS-CoV-2 Detected | SARS-CoV-2 Detected | 16879824 | 15895954 | 5.42 | 93.21 | 15034264 | 98.55 | 526674561 | 17612.76665 | 99.96655854 | 10 | 29893 | 51 | 76 | 12 | 129 |
| CS0537_1 | CS0537_2 | Positive | Positive | SARS-CoV-2 Detected | SARS-CoV-2 Detected | 17737571 | 16645780 | 0.9 | 97.79 | 16495243 | 98.69 | 578638743 | 19350.5248 | 99.97659098 | 7 | 29896 | 51 | 251 | 12 | 312 |
| CS0536_1 | CS0536_2 | Positive | Positive | SARS-CoV-2 Detected | SARS-CoV-2 Detected | 17579963 | 16466263 | 0.85 | 97.66 | 16325680 | 98.5 | 571621499 | 19115.85791 | 99.93980537 | 18 | 29885 | 74 | 240 | 11 | 319 |
| CS0535_1 | CS0535_2 | Positive | Positive | SARS-CoV-2 Detected | SARS-CoV-2 Detected | 15300331 | 14133495 | 6.11 | 92.6 | 13269910 | 98.63 | 465018803 | 15550.90804 | 99.99665585 | 1 | 29902 | 44 | 436 | 12 | 295 |
| CS0534_1 | CS0534_2 | Positive | Positive | SARS-CoV-2 Detected | SARS-CoV-2 Detected | 25164430 | 23582985 | 0.9 | 98.26 | 23369824 | 99.16 | 823617472 | 27542.97134 | 100 | 0 | 29903 | 46 | 85 | 12 | 68 |
| CS0551_1 | CS0551_2 | Positive | Positive | SARS-CoV-2 Detected | SARS-CoV-2 Detected | 16531650 | 15499827 | 5 | 93.83 | 14725197 | 98.77 | 516972692 | 17288.32197 | 99.82944855 | 51 | 29852 | 80 | 323 | 15 | 287 |
| [illegible] | | | | | | | | | | | | | | | | | | | | |

| | In - Depth Custom Pipeline (Replicate 1) | | | | | | | | | | | | | | | |
|---|---|---|---|---|---|---|---|---|---|---|---|---|---|---|---|---|
| Sample IDs Replicate 1 | Total Reads_Replicate1 | Trimmed Reads q30-30bp_Replicate1 | Kraken2 coronavirus %_Replicate1 | Hisat2 Human % _Replicate1 | Hisat2 Covidall % _Replicate1 | Hisat2 Unmapped Covid Read#_Replicate1 | Hisat2 Unmapped Covid %_Replicate1 | BasePairescovered_Replicate1 | X coverage _Replicate1 | Genome coverage_Replicate1 | # bases with 0 coverage_Replicate1 | # bases with >0 coverage_Replicate1 | # bases with <10x coverage_Replicate1 | # bases with <100x coverage_Replicate1 | VARSCAN Variant Count_Replicate1 | N's_Replicate1 |
| CS0531_1 | 9227700 | 8702460 | 0.54 | 1.12 | 98.08 | 8605386 | 99.19 | 303345230 | 10144.30759 | 99.93980537 | 18 | 29885 | 71 | 110 | 13 | 58 |
| CS0530_1 | 7083108 | 6666395 | 0.54 | 4.3 | 94.63 | 6379569 | 98.88 | 224250778 | 7499.273585 | 99.84616928 | 46 | 29857 | 52 | 472 | 15 | 231 |
| CS0529_1 | 8561478 | 8080890 | 0.56 | 4.29 | 94.65 | 7734194 | 98.89 | 271879245 | 9092.039093 | 99.83613684 | 49 | 29854 | 71 | 562 | 15 | 287 |
| CS0533_1 | 8195160 | 7741017 | 0.63 | 1.1 | 97.94 | 7655629 | 99.03 | 269581080 | 9015.185098 | 99.76256563 | 71 | 29832 | 111 | 917 | 13 | 323 |
| CS0545_1 | 9570945 | 8971591 | 0.59 | 1.05 | 98.03 | 8877643 | 99.06 | 312526290 | 10451.33565 | 99.94314952 | 17 | 29886 | 63 | 180 | 11 | 112 |
| CS0532_1 | 9623488 | 9027824 | 0.5 | 7.39 | 91.11 | 8360737 | 98.38 | 292354092 | 9776.747885 | 99.95318195 | 14 | 29889 | 59 | 243 | 14 | 302 |
| CS0544_1 | 10051772 | 9506119 | 0.58 | 0.94 | 98.2 | 9417024 | 99.13 | 331917289 | 11099.79898 | 99.95987025 | 12 | 29891 | 70 | 567 | 11 | 321 |
| CS0543_1 | 9758391 | 9274190 | 0.55 | 4.7 | 94.24 | 8838312 | 98.89 | 310759485 | 10392.25111 | 99.78597465 | 64 | 29839 | 82 | 517 | 15 | 309 |
| CS0542_1 | 7248976 | 6858700 | 0.52 | 8.4 | 89.87 | 6282236 | 98.12 | 219121755 | 7327.751563 | 99.96321439 | 11 | 29892 | 55 | 932 | 14 | 435 |
| CS0541_1 | 10211538 | 9525628 | 0.54 | 3.01 | 95.94 | 9239217 | 98.91 | 324752808 | 10860.20827 | 99.85620172 | 43 | 29860 | 52 | 221 | 15 | 176 |
| CS0540_1 | 9132247 | 8627809 | 0.55 | 3.32 | 95.74 | 8341717 | 99.02 | 293641172 | 9819.78972 | 99.94649366 | 16 | 29887 | 58 | 330 | 12 | 309 |
| CS0539_1 | 8928587 | 8422471 | 0.56 | 1.44 | 97.77 | 8301302 | 99.19 | 292750428 | 9790.00194 | 99.84951343 | 45 | 29858 | 79 | 175 | 10 | 293 |
| CS0538_1 | 8449838 | 8002331 | 0.57 | 5.47 | 93.15 | 7564696 | 98.54 | 264999422 | 8861.967762 | 99.84616928 | 46 | 29857 | 56 | 91 | 12 | 236 |
| CS0537_1 | 8890929 | 8395117 | 0.57 | 0.9 | 97.79 | 8319425 | 98.68 | 291850025 | 9759.891148 | 99.96990269 | 9 | 29894 | 72 | 446 | 12 | 331 |
| CS0536_1 | 8811763 | 8307149 | 0.58 | 0.86 | 97.65 | 8235646 | 98.49 | 288366492 | 9643.396716 | 99.93980537 | 18 | 29885 | 89 | 870 | 11 | 341 |
| CS0535_1 | 7670372 | 7143534 | 0.53 | 6.15 | 92.56 | 6704270 | 98.62 | 234946158 | 7856.942715 | 99.90636391 | 28 | 29875 | 54 | 718 | 12 | 283 |
| CS0534_1 | 12583680 | 11868828 | 0.56 | 0.91 | 98.25 | 11761142 | 99.15 | 414515586 | 13862.00669 | 99.86289001 | 41 | 29862 | 52 | 95 | 13 | 149 |
| CS0551_1 | 8286688 | 7819397 | 0.56 | 5.02 | 93.8 | 7426489 | 98.76 | 260742208 | 8719.600308 | 99.82944855 | 51 | 29852 | 86 | 685 | 14 | 303 |
| [illegible] | | | | | | | | | | | | | | | | |

| | In - Depth Custom Pipeline (Replicate 2) | | | | | | | | | | | | | | |
|---|---|---|---|---|---|---|---|---|---|---|---|---|---|---|---|
| Sample IDs Replicate 2 | Total Reads_Replicate2 | Trimmed Reads q30-30bp_Replicate2 | Hisat2 Human % _Replicate2 | Hisat2 Covidall % _Replicate2 | Hisat2 Unmapped Covid Read#_Replicate2 | Hisat2 Unmapped Covid %_Replicate2 | BasePairescovered_Replicate2 | X coverage _Replicate2 | Genome coverage_Replicate2 | # bases with 0 coverage_Replicate2 | # bases with >0 coverage_Replicate2 | # bases with <10x coverage_Replicate2 | # bases with <100x coverage_Replicate2 | VARSCAN Variant Count_Replicate2 | N's_Replicate2 |
| CS0531_2 | 9124885 | 8492501 | 1.11 | 98.08 | 8398317 | 99.18 | 295960050 | 9897.336388 | 99.94649366 | 16 | 29887 | 70 | 134 | 13 | 116 |
| CS0530_2 | 7027432 | 6523869 | 4.25 | 94.68 | 6246688 | 98.88 | 219528086 | 7341.339866 | 99.86289001 | 41 | 29862 | 52 | 494 | 14 | 271 |
| CS0529_2 | 8503025 | 7922914 | 4.24 | 94.69 | 7587053 | 98.88 | 266644678 | 8916.987526 | 99.85954586 | 42 | 29861 | 64 | 553 | 15 | 281 |
| CS0533_2 | 8167969 | 7628223 | 1.09 | 97.97 | 7545317 | 99.05 | 265706286 | 8885.606327 | 99.78263051 | 65 | 29838 | 86 | 400 | 14 | 427 |
| CS0545_2 | 9481183 | 8761915 | 1.04 | 98.03 | 8670766 | 99.06 | 305167094 | 10205.23339 | 99.83948099 | 48 | 29855 | 64 | 167 | 12 | 79 |
| CS0532_2 | 9593313 | 8868074 | 7.29 | 91.23 | 8221760 | 98.4 | 287485997 | 9613.951677 | 99.9565261 | 13 | 29890 | 59 | 264 | 12 | 315 |
| CS0544_2 | 10026104 | 9364779 | 0.93 | 98.22 | 9277565 | 99.14 | 326956731 | 10933.91068 | 99.83948099 | 48 | 29855 | 78 | 546 | 11 | 294 |
| CS0543_2 | 9727621 | 9142629 | 4.65 | 94.29 | 8717514 | 98.89 | 306463155 | 10248.57556 | 99.83948099 | 48 | 29855 | 76 | 498 | 15 | 302 |
| CS0542_2 | 7214723 | 6745322 | 8.26 | 90.04 | 6187897 | 98.16 | 215881175 | 7219.381835 | 99.85285757 | 44 | 29859 | 54 | 303 | 15 | 374 |
| CS0541_2 | 10123540 | 9298071 | 2.98 | 95.97 | 9021154 | 98.91 | 317018877 | 10601.57432 | 99.95318195 | 14 | 29889 | 63 | 186 | 15 | 225 |
| CS0540_2 | 9035247 | 8425463 | 3.28 | 95.77 | 8148805 | 99.02 | 286816010 | 9591.546333 | 99.82944855 | 51 | 29852 | 59 | 574 | 12 | 264 |
| CS0539_2 | 8899058 | 8289399 | 1.42 | 97.78 | 8171425 | 99.19 | 288108101 | 9634.755744 | 99.81272782 | 56 | 29847 | 77 | 233 | 10 | 293 |
| CS0538_2 | 8429986 | 7893623 | 5.37 | 93.26 | 7469614 | 98.55 | 261674350 | 8750.772498 | 99.96321439 | 11 | 29892 | 53 | 119 | 12 | 209 |
| CS0537_2 | 8846642 | 8250663 | 0.91 | 97.79 | 8175800 | 98.69 | 286788422 | 9590.62375 | 99.83948099 | 48 | 29855 | 60 | 481 | 12 | 268 |
| CS0536_2 | 8768200 | 8159114 | 0.84 | 97.67 | 8090175 | 98.51 | 283254782 | 9472.453667 | 99.92308464 | 23 | 29880 | 80 | 440 | 11 | 349 |
| CS0535_2 | 7629959 | 6989961 | 6.07 | 92.65 | 6565956 | 98.63 | 230072232 | 7693.95151 | 99.99331171 | 2 | 29901 | 51 | 693 | 11 | 311 |
| CS0534_2 | 12580750 | 11714157 | 0.9 | 98.27 | 11608682 | 99.16 | 409101607 | 13680.95532 | 100 | 0 | 29903 | 61 | 88 | 13 | 214 |
| CS0551_2 | 8244962 | 7680430 | 4.96 | 93.86 | 7299097 | 98.77 | 256230474 | 8568.721332 | 99.7893188 | 63 | 29840 | 93 | 608 | 15 | 304 |
| [illegible] | | | | | | | | | | | | | | | |

| | | RT-PCR | | DRAGEN COVIDSeq Test Pipeline | | In - Depth Custom Pipeline (Merged) | | | | | | | | | | | | | | |
|---|---|---|---|---|---|---|---|---|---|---|---|---|---|---|---|---|---|---|---|---|
| Sample IDs Replicate 1 | Sample IDs Replicate 2 | RT (Pos/Neg) replicate1 | RT (Pos/Neg) replicate 2 | DRAGEN SARS-CoV-2 Detection Replicate 1 | DRAGEN SARS-CoV-2 Detection Replicate 2 | Total Reads_merged | Trimmed Reads q30-30bp_merged | Hisat2 Human % _merged | Hisat2 Covidall % _merged | Hisat2 Unmapped Covid Read#_merged | Hisat2 Unmapped Covid %_merged | BasePairscovered_merged | X coverage _merged | Genome coverage_merged | # bases with 0 coverage_merged | # bases with >0 coverage_merged | # bases with <10x coverage_merged | # bases with <100x coverage_merged | VARSCAN Variant Count_merged | N's_merged |
| CS0036_1 | CS0036_2 | Positive | Positive | SARS-CoV-2 Detected | SARS-CoV-2 Detected | 19610113 | 18531332 | 0.01 | 99.28 | 18529153 | 99.3 | 654195859 | 21877.26512 | 99.77594221 | 67 | 29836 | 82 | 479 | 13 | 292 |
| CS0279_1 | CS0279_2 | Positive | Positive | SARS-CoV-2 Detected | SARS-CoV-2 Detected | 22713385 | 21446803 | 1.93 | 97.07 | 21033550 | 98.97 | 740178330 | 24752.64455 | 99.98996756 | 3 | 29900 | 50 | 350 | 14 | 258 |
| CS0280_1 | CS0280_2 | Positive | Positive | SARS-CoV-2 Detected | SARS-CoV-2 Detected | 11836612 | 11170908 | 69.51 | 25.87 | 3405751 | 84.85 | 102828343 | 3438.729994 | 87.84068488 | 3636 | 26267 | 23840 | 25585 | 8 | 22152 |
| CS0724_1 | CS0724_2 | Inconclusive | Inconclusive | SARS-CoV-2 Detected | SARS-CoV-2 Detected | 17673223 | 16508247 | 37.14 | 59.44 | 10377623 | 94.56 | 349034127 | 11672.21105 | 96.42845199 | 1068 | 28835 | 14113 | 16243 | 8 | 14153 |
| CS0281_1 | CS0281_2 | Positive | Positive | SARS-CoV-2 Detected | SARS-CoV-2 Detected | 17976729 | 16954611 | 0.19 | 98.31 | 16922135 | 98.5 | 592600727 | 19817.43394 | 99.8327927 | 50 | 29853 | 71 | 275 | 14 | 291 |
| CS0725_1 | CS0725_2 | Inconclusive | Inconclusive | SARS-CoV-2 Not Detected | SARS-CoV-2 Not Detected | 23651483 | 22319401 | 94.91 | 0.08 | 1135164 | 1.53 | 618761 | 20.69227168 | 5.654951008 | 28212 | 1691 | 29086 | 29131 | 0 | 29103 |
| CS0726_1 | CS0726_2 | Inconclusive | Inconclusive | SARS-CoV-2 Detected | SARS-CoV-2 Detected | 17094792 | 16128502 | 73.81 | 14.56 | 4224636 | 55.59 | 83579185 | 2795.010032 | 37.63836404 | 18648 | 11255 | 19489 | 20806 | 8 | 19491 |
| CS0282_1 | CS0282_2 | Positive | Positive | SARS-CoV-2 Detected | SARS-CoV-2 Detected | 17313673 | 16090024 | 1.86 | 96.84 | 15791248 | 98.67 | 553834724 | 18521.04217 | 99.87292245 | 38 | 29865 | 109 | 830 | 12 | 352 |
| CS0523_1 | CS0523_2 | Pan Sarbeco | Pan Sarbeco | SARS-CoV-2 Not Detected | SARS-CoV-2 Not Detected | 6890935 | 6443455 | 98.3 | 0.53 | 109574 | 31.45 | 1216695 | 40.68805805 | 7.370497943 | 27699 | 2204 | 29463 | 29497 | 1 | 29457 |
| CS0037_1 | CS0037_2 | Positive | Positive | SARS-CoV-2 Detected | SARS-CoV-2 Detected | 22511342 | 21031966 | 0.08 | 35.31 | 21015731 | 35.33 | 263933228 | 8826.312678 | 99.84616928 | 46 | 29857 | 198 | 1613 | 6 | 547 |
| CS0038_1 | CS0038_2 | Positive | Positive | SARS-CoV-2 Detected | SARS-CoV-2 Detected | 17844959 | 16720627 | 2.44 | 96.52 | 16312285 | 98.94 | 573768505 | 19187.65692 | 99.73915661 | 78 | 29825 | 269 | 1046 | 11 | 556 |
| CS0039_1 | CS0039_2 | Positive | Positive | SARS-CoV-2 Detected | SARS-CoV-2 Detected | 15467905 | 14592754 | 0.77 | 98.29 | 14479740 | 99.06 | 509885087 | 17051.30211 | 99.86289001 | 41 | 29862 | 50 | 551 | 12 | 82 |
| CS0286_1 | CS0286_2 | Positive | Positive | SARS-CoV-2 Detected | SARS-CoV-2 Detected | 15286579 | 14188010 | 90.69 | 1.58 | 1320671 | 16.96 | 7956276 | 266.0694914 | 32.77597565 | 20102 | 9801 | 22395 | 24561 | 7 | 22030 |
| CS0727_1 | CS0727_2 | Positive | Positive | SARS-CoV-2 Detected | SARS-CoV-2 Detected | 19224302 | 18104882 | 89.47 | 4.04 | 1906491 | 38.33 | 26005097 | 869.6484299 | 19.69367622 | 24014 | 5889 | 25336 | 26235 | 2 | 25307 |
| CS0287_1 | CS0287_2 | Positive | Positive | SARS-CoV-2 Detected | SARS-CoV-2 Detected | 15101207 | 14243393 | 5.84 | 93.17 | 13412076 | 98.94 | 471744791 | 15775.8349 | 99.85285757 | 44 | 29859 | 58 | 88 | 11 | 94 |
| CS0040_1 | CS0040_2 | Positive | Positive | SARS-CoV-2 Detected | SARS-CoV-2 Detected | 10203051 | 9651882 | 59.65 | 31.97 | 3894268 | 79.24 | 109812460 | 3672.289068 | 45.34327659 | 16344 | 13559 | 17691 | 18946 | 6 | 17352 |
| CS0288_1 | CS0288_2 | Positive | Positive | SARS-CoV-2 Detected | SARS-CoV-2 Detected | 15855572 | 14952756 | 5.29 | 93.61 | 14162260 | 98.84 | 497649790 | 16642.13591 | 99.80603953 | 58 | 29845 | 74 | 90 | 16 | 217 |
| CS0289_1 | CS0289_2 | Positive | Positive | SARS-CoV-2 Detected | SARS-CoV-2 Detected | 16597900 | 15664348 | 16.96 | 81.02 | 13007748 | 97.56 | 451350779 | 15093.82935 | 99.26428786 | 220 | 29683 | 558 | 2349 | 14 | 600 |
| [illegible] | [illegible] | [illegible] | [illegible] | [illegible] | [illegible] | [illegible] | [illegible] | [illegible] | [illegible] | [illegible] | [illegible] | [illegible] | [illegible] | [illegible] | [illegible] | [illegible] | [illegible] | [illegible] | [illegible] | [illegible] |

| | In - Depth Custom Pipeline (Replicate 1) | | | | | | | | | | | | | | | |
|---|---|---|---|---|---|---|---|---|---|---|---|---|---|---|---|---|
| Sample IDs Replicate 1 | Total Reads_Replicate1 | Trimmed Reads q30-30bp_Replicate1 | Kraken2 coronavirus %_Replicate1 | Hisat2 Human % _Replicate1 | Hisat2 Covidall % _Replicate1 | Hisat2 Unmapped Covid Read#_Replicate1 | Hisat2 Unmapped Covid %_Replicate1 | BasePairscovered_Replicate1 | X coverage _Replicate1 | Genome coverage_Replicate1 | # bases with 0 coverage_Replicate1 | # bases with >0 coverage_Replicate1 | # bases with <10x coverage_Replicate1 | # bases with <100x coverage_Replicate1 | VARSCAN Variant Count_Replicate1 | N's_Replicate1 |
| CS0036_1 | 9827918 | 9323575 | 0.63 | 0.01 | 99.29 | 9322467 | 99.31 | 329206234 | 11009.13734 | 99.77594221 | 67 | 29836 | 89 | 641 | 13 | 312 |
| CS0279_1 | 11363497 | 10775703 | 0.65 | 1.93 | 97.07 | 10567579 | 98.98 | 371944247 | 12438.35893 | 99.9565261 | 13 | 29890 | 67 | 804 | 14 | 289 |
| CS0280_1 | 5955397 | 5640253 | 0.21 | 69.47 | 25.85 | 1721693 | 84.68 | 51883114 | 1735.047119 | 22.5662977 | 23155 | 6748 | 24270 | 25838 | 7 | 22431 |
| CS0724_1 | 8876822 | 8345614 | 0.59 | 37.14 | 59.34 | 5246255 | 94.39 | 176161522 | 5891.098619 | 55.61983747 | 13271 | 16632 | 15111 | 17566 | 7 | 14038 |
| CS0281_1 | 9014238 | 8538114 | 0.65 | 0.19 | 98.33 | 8521825 | 98.51 | 298501911 | 9982.339932 | 99.8327927 | 50 | 29853 | 73 | 398 | 14 | 291 |
| CS0725_1 | 11895106 | 11284772 | 0 | 94.88 | 0.08 | 577264 | 1.51 | 309130 | 10.33775875 | 4.093234792 | 28679 | 1224 | 29113 | 29297 | 0 | 29112 |
| CS0726_1 | 8589161 | 8148078 | 0.1 | 73.8 | 14.49 | 2134407 | 55.32 | 42030229 | 1405.552252 | 36.58830218 | 18962 | 10941 | 19677 | 21326 | 8 | 19636 |
| CS0282_1 | 8664377 | 8094378 | 0.67 | 1.86 | 96.84 | 7944115 | 98.68 | 278667322 | 9319.042303 | 99.76256563 | 71 | 29832 | 235 | 1823 | 12 | 368 |
| CS0523_1 | 3465844 | 3261191 | 0 | 98.3 | 0.54 | 55527 | 31.49 | 616934 | 20.63117413 | 1.625254991 | 29417 | 486 | 29466 | 29497 | 1 | 24032 |
| CS0037_1 | 11314435 | 10626972 | 0.23 | 0.08 | 35.35 | 10618739 | 35.38 | 133553710 | 4466.231147 | 99.8261044 | 52 | 29851 | 673 | 3076 | 6 | 702 |
| CS0038_1 | 8906634 | 8385381 | 0.66 | 2.45 | 96.53 | 8180164 | 98.95 | 287791760 | 9624.176838 | 99.69902685 | 90 | 29813 | 332 | 2186 | 11 | 912 |
| CS0039_1 | 7787632 | 7376760 | 0.6 | 0.78 | 98.3 | 7319347 | 99.07 | 257789857 | 8620.869378 | 99.85620172 | 43 | 29860 | 53 | 817 | 12 | 218 |
| CS0286_1 | 7708675 | 7187096 | 0.01 | 90.67 | 1.57 | 670344 | 16.82 | 4005108 | 133.9366619 | 31.27779821 | 20550 | 9353 | 22972 | 25426 | 7 | 22818 |
| CS0727_1 | 9694055 | 9180476 | 0.06 | 89.38 | 4.02 | 975276 | 37.88 | 13148638 | 439.7096612 | 18.64027021 | 24329 | 5574 | 25503 | 26354 | 1 | 25461 |
| CS0287_1 | 7584878 | 7182018 | 0.56 | 5.87 | 93.14 | 6760349 | 98.95 | 237817640 | 7952.969267 | 99.84616928 | 46 | 29857 | 69 | 92 | 13 | 252 |
| CS0040_1 | 5128036 | 4867698 | 0.43 | 59.69 | 31.95 | 1962062 | 79.26 | 55342191 | 1850.723707 | 44.38350667 | 16631 | 13272 | 18110 | 19614 | 5 | 17813 |
| CS0288_1 | 7938173 | 7518412 | 0.59 | 5.28 | 93.62 | 7121479 | 98.84 | 250278762 | 8369.687389 | 99.79935124 | 60 | 29843 | 85 | 244 | 16 | 290 |
| CS0289_1 | 8341367 | 7902148 | 0.64 | 16.96 | 81.02 | 6561773 | 97.57 | 227711025 | 7614.989299 | 98.99341203 | 301 | 29602 | 861 | 3415 | 14 | 935 |
| [illegible] | [illegible] | [illegible] | [illegible] | [illegible] | [illegible] | [illegible] | [illegible] | [illegible] | [illegible] | [illegible] | [illegible] | [illegible] | [illegible] | [illegible] | [illegible] | [illegible] |

| | In - Depth Custom Pipeline (Replicate 2) | | | | | | | | | | | | | | |
|---|---|---|---|---|---|---|---|---|---|---|---|---|---|---|---|
| Sample IDs Replicate 2 | Total Reads_Replicate2 | Trimmed Reads q30-30bp_Replicate2 | Hisat2 Human % _Replicate2 | Hisat2 Covidall % _Replicate2 | Hisat2 Unmapped Covid Read#_Replicate2 | Hisat2 Unmapped Covid %_Replicate2 | BasePairscovered_Replicate2 | X coverage _Replicate2 | Genome coverage_Replicate2 | # bases with 0 coverage_Replicate2 | # bases with >0 coverage_Replicate2 | # bases with <10x coverage_Replicate2 | # bases with <100x coverage_Replicate2 | VARSCAN Variant Count_Replicate2 | N's_Replicate2 |
| CS0036_2 | 9782195 | 9207757 | 0.01 | 99.27 | 9206686 | 99.29 | 324989637 | 10868.12818 | 99.76925392 | 69 | 29834 | 87 | 892 | 13 | 303 |
| CS0279_2 | 11349888 | 10671100 | 1.92 | 97.06 | 10465971 | 98.97 | 368234083 | 12314.28562 | 99.92308464 | 23 | 29880 | 56 | 488 | 15 | 290 |
| CS0280_2 | 5881215 | 5530655 | 69.48 | 25.89 | 1687837 | 84.83 | 50945229 | 1703.682875 | 87.35912785 | 3780 | 26123 | 24220 | 25891 | 7 | 21199 |
| CS0724_2 | 8796401 | 8162633 | 36.46 | 59.55 | 5186238 | 93.73 | 172872605 | 5781.11243 | 96.41841956 | 1071 | 28832 | 14911 | 17667 | 7 | 14911 |
| CS0281_2 | 8962491 | 8416497 | 0.19 | 98.29 | 8400310 | 98.48 | 294098825 | 9835.094305 | 99.80938367 | 57 | 29846 | 82 | 403 | 14 | 293 |
| CS0725_2 | 11756377 | 11034629 | 94.94 | 0.08 | 558044 | 1.56 | 309631 | 10.35451293 | 4.534662074 | 28547 | 1356 | 29113 | 29262 | 0 | 29106 |
| CS0726_2 | 8505631 | 7980424 | 73.81 | 14.63 | 2090237 | 55.85 | 41548956 | 1389.45778 | 36.65852924 | 18941 | 10962 | 19614 | 21295 | 8 | 19582 |
| CS0282_2 | 8649296 | 7995646 | 1.86 | 96.83 | 7847217 | 98.66 | 275167398 | 9201.999732 | 99.85285757 | 44 | 29859 | 220 | 1900 | 12 | 446 |
| CS0523_2 | 3425091 | 3182264 | 98.3 | 0.53 | 54047 | 31.41 | 599761 | 20.05688392 | 7.246764539 | 27736 | 2167 | 29467 | 29497 | 1 | 29467 |
| CS0037_2 | 11196907 | 10404994 | 0.08 | 35.26 | 10397003 | 35.29 | 130379528 | 4360.081865 | 99.72912417 | 81 | 29822 | 708 | 2845 | 6 | 713 |
| CS0038_2 | 8938325 | 8335246 | 2.44 | 96.51 | 8132126 | 98.92 | 285976709 | 9563.478882 | 99.67227368 | 98 | 29805 | 347 | 1784 | 11 | 941 |
| CS0039_2 | 7680273 | 7215994 | 0.77 | 98.28 | 7160665 | 99.04 | 252095276 | 8430.434271 | 99.85954586 | 42 | 29861 | 52 | 817 | 12 | 224 |
| CS0286_2 | 7577904 | 7000914 | 90.25 | 1.59 | 682901 | 16.29 | 3951168 | 132.1328295 | 31.18081798 | 20579 | 9324 | 22892 | 25394 | 6 | 22793 |
| CS0727_2 | 9530247 | 8924406 | 89.23 | 4.05 | 961344 | 37.58 | 12856459 | 429.9387687 | 17.19894325 | 24760 | 5143 | 25533 | 26391 | 2 | 25119 |
| CS0287_2 | 7516329 | 7061375 | 5.77 | 93.2 | 6653598 | 98.91 | 233926652 | 7822.848945 | 99.85285757 | 44 | 29859 | 68 | 351 | 13 | 194 |
| CS0040_2 | 5075015 | 4784184 | 59.65 | 32 | 1930261 | 79.31 | 54470269 | 1821.565361 | 43.73808648 | 16824 | 13079 | 18136 | 19735 | 6 | 17810 |
| CS0288_2 | 7917399 | 7434344 | 5.28 | 93.6 | 7041747 | 98.82 | 247371946 | 8272.479216 | 99.80603953 | 58 | 29845 | 80 | 482 | 16 | 296 |
| CS0289_2 | 8256533 | 7762200 | 16.91 | 81.02 | 6449643 | 97.5 | 223639763 | 7478.84035 | 99.05026252 | 284 | 29619 | 1134 | 4407 | 14 | 951 |
| [illegible] | [illegible] | [illegible] | [illegible] | [illegible] | [illegible] | [illegible] | [illegible] | [illegible] | [illegible] | [illegible] | [illegible] | [illegible] | [illegible] | [illegible] | [illegible] |

| Sample IDs Replicate 1 | Sample IDs Replicate 2 | RT-PCR: RT (Pos/Neg) replicate1 | RT (Pos/Neg) replicate 2 | DRAGEN COVIDSeq Test Pipeline: DRAGEN SARS-CoV-2 Detection Replicate 1 | DRAGEN SARS-CoV-2 Detection Replicate 2 | In - Depth Custom Pipeline (Merged): Total Reads_merged | Trimmed Reads q30-30bp_merged | Hisat2 Human % _merged | Hisat2 Covidall % _merged | Hisat2 Unmapped Covid Read#_merged | Hisat2 Unmapped Covid %_merged | BasePairescovered_merged | X coverage _merged | Genome coverage_merged | # bases with 0 coverage_merged | # bases with >0 coverage_merged | # bases with <10x coverage_merged | # bases with <100x coverage_merged | VARSCAN Variant Count_merged | N's_merged |
|---|---|---|---|---|---|---|---|---|---|---|---|---|---|---|---|---|---|---|---|---|
| CS0169_1 | CS0169_2 | Positive | Positive | SARS-CoV-2 Detected | SARS-CoV-2 Detected | 17106701 | 15999130 | 25.14 | 72.12 | 11977706 | 96.33 | 410138680 | 13715.63656 | 99.86623416 | 40 | 29863 | 267 | 876 | 11 | 551 |
| CS0170_1 | CS0170_2 | Positive | Positive | SARS-CoV-2 Detected | SARS-CoV-2 Detected | 18186377 | 17160358 | 22.3 | 74.93 | 13333215 | 96.44 | 457142079 | 15287.49888 | 99.94314952 | 17 | 29886 | 79 | 137 | 12 | 339 |
| CS0171_1 | CS0171_2 | Positive | Positive | SARS-CoV-2 Detected | SARS-CoV-2 Detected | 20802936 | 19354938 | 14.71 | 83.36 | 16508099 | 97.73 | 573319608 | 19172.64515 | 99.96655854 | 10 | 29893 | 72 | 211 | 16 | 286 |
| CS0634_1 | CS0634_2 | Positive | Positive | SARS-CoV-2 Detected | SARS-CoV-2 Detected | 20894380 | 19726581 | 86.6 | 7.11 | 2643399 | 53.03 | 49862528 | 1667.475772 | 82.29274655 | 5295 | 24608 | 7935 | 11951 | 15 | 7687 |
| CS0172_1 | CS0172_2 | Positive | Positive | SARS-CoV-2 Detected | SARS-CoV-2 Detected | 18079813 | 17124816 | 5.67 | 93.16 | 16153350 | 98.77 | 567252335 | 18969.74668 | 99.97324683 | 8 | 29895 | 71 | 232 | 13 | 282 |
| CS0173_1 | CS0173_2 | Positive | Positive | SARS-CoV-2 Detected | SARS-CoV-2 Detected | 15501956 | 14578844 | 20.24 | 74.24 | 11627360 | 93.08 | 384774999 | 12867.43802 | 99.28435274 | 214 | 29689 | 360 | 927 | 12 | 844 |
| CS0174_1 | CS0174_2 | Positive | Positive | SARS-CoV-2 Detected | SARS-CoV-2 Detected | 16263368 | 15197179 | 29.34 | 64.05 | 10738776 | 90.64 | 345979895 | 11570.07307 | 99.82276026 | 53 | 29850 | 355 | 1042 | 17 | 583 |
| CS0572_1 | CS0572_2 | Positive | Positive | SARS-CoV-2 Detected | SARS-CoV-2 Detected | 17761726 | 15990317 | 87.33 | 2.71 | 2026391 | 21.41 | 15405581 | 515.1851319 | 43.9721767 | 16754 | 13149 | 17739 | 19549 | 9 | 17662 |
| CS0175_1 | CS0175_2 | Positive | Positive | SARS-CoV-2 Detected | SARS-CoV-2 Detected | 20695812 | 19421234 | 1.32 | 97.78 | 19165750 | 99.09 | 674810342 | 22566.64355 | 99.86623416 | 40 | 29863 | 46 | 62 | 13 | 8 |
| CS0635_1 | CS0635_2 | Positive | Positive | SARS-CoV-2 Detected | SARS-CoV-2 Detected | 27501943 | 25926776 | 59.97 | 35.27 | 10379486 | 88.1 | 325218156 | 10875.77019 | 99.84616928 | 46 | 29857 | 131 | 2609 | 15 | 1368 |
| CS0405_1 | CS0405_2 | Pan Sarbeco | Pan Sarbeco | SARS-CoV-2 Not Detected | SARS-CoV-2 Not Detected | 13893480 | 13037813 | 96.68 | 0.01 | 432484 | 0.35 | 53233 | 1.780189279 | 8.146339832 | 27467 | 2436 | 28588 | 29876 | 2 | 28543 |
| CS0406_1 | CS0406_2 | Pan Sarbeco | Pan Sarbeco | SARS-CoV-2 Not Detected | SARS-CoV-2 Not Detected | 18349251 | 17288917 | 95.34 | 0.07 | 806148 | 1.42 | 408321 | 13.65485068 | 7.179881617 | 27756 | 2147 | 28767 | 29098 | 0 | 28764 |
| CS0636_1 | CS0636_2 | Positive | Positive | SARS-CoV-2 Detected | SARS-CoV-2 Detected | 19481569 | 18355194 | 86.47 | 8.45 | 2482681 | 62.47 | 55114928 | 1843.1237 | 83.57689864 | 4911 | 24992 | 11012 | 13855 | 10 | 10888 |
| CS0176_1 | CS0176_2 | Positive | Positive | SARS-CoV-2 Detected | SARS-CoV-2 Detected | 12941379 | 12224521 | 33.1 | 62.58 | 8177765 | 93.55 | 271990674 | 9095.765442 | 99.4850015 | 154 | 29749 | 426 | 946 | 12 | 845 |
| CS0637_1 | CS0637_2 | Inconclusive | Inconclusive | SARS-CoV-2 Detected | SARS-CoV-2 Detected | 14647798 | 13744693 | 93.87 | 1.16 | 843126 | 18.98 | 5689268 | 190.2574324 | 9.226499013 | 27144 | 2759 | 28022 | 28395 | 0 | 28014 |
| CS0177_1 | CS0177_2 | Positive | Positive | SARS-CoV-2 Detected | SARS-CoV-2 Detected | 17541381 | 16532526 | 30.78 | 60.95 | 11442990 | 88.06 | 358259794 | 11980.73083 | 99.87961074 | 36 | 29867 | 68 | 399 | 15 | 560 |
| CS0178_1 | CS0178_2 | Positive | Positive | SARS-CoV-2 Detected | SARS-CoV-2 Detected | 18369317 | 17153727 | 38.01 | 57.14 | 10634209 | 92.18 | 348430758 | 11652.03351 | 99.86289001 | 41 | 29862 | 243 | 647 | 11 | 747 |
| CS0638_1 | CS0638_2 | Positive | Positive | SARS-CoV-2 Detected | SARS-CoV-2 Detected | 17219549 | 16222567 | 87.36 | 5.91 | 2049779 | 46.81 | 34132517 | 1141.441227 | 87.67013343 | 3687 | 26216 | 12181 | 17037 | 20 | 11999 |

| Sample IDs Replicate 1 | In - Depth Custom Pipeline (Replicate 1): Total Reads_Replicate1 | Trimmed Reads q30-30bp_Replicate1 | Kraken2 coronavirus %_Replicate1 | Hisat2 Human % _Replicate1 | Hisat2 Covidall % _Replicate1 | Hisat2 Unmapped Covid Read#_Replicate1 | Hisat2 Unmapped Covid %_Replicate1 | BasePairescovered_Replicate1 | X coverage _Replicate1 | Genome coverage_Replicate1 | # bases with 0 coverage_Replicate1 | # bases with >0 coverage_Replicate1 | # bases with <10x coverage_Replicate1 | # bases with <100x coverage_Replicate1 | VARSCAN Variant Count_Replicate1 | N's_Replicate1 |
|---|---|---|---|---|---|---|---|---|---|---|---|---|---|---|---|---|
| CS0169_1 | 8551635 | 8034941 | 0.49 | 25.12 | 72.12 | 6016369 | 96.32 | 205992751 | 6888.698492 | 99.70571515 | 88 | 29815 | 578 | 1784 | 12 | 864 |
| CS0170_1 | 9093586 | 8613614 | 0.49 | 22.32 | 74.9 | 6690913 | 96.42 | 229378952 | 7670.767214 | 99.88295489 | 35 | 29868 | 85 | 381 | 10 | 386 |
| CS0171_1 | 10422576 | 9751219 | 0.5 | 14.72 | 83.34 | 8315788 | 97.72 | 288796238 | 9657.76805 | 99.94983781 | 15 | 29888 | 79 | 462 | 16 | 485 |
| CS0634_1 | 10544326 | 10009901 | 0.07 | 86.52 | 7.07 | 1349072 | 52.43 | 25157094 | 841.2899709 | 80.56716717 | 5811 | 24092 | 9014 | 14722 | 13 | 8627 |
| CS0172_1 | 9074924 | 8630511 | 0.57 | 5.67 | 93.17 | 8141060 | 98.77 | 285921721 | 9561.640003 | 99.86289001 | 41 | 29862 | 80 | 758 | 13 | 291 |
| CS0173_1 | 7790329 | 7357527 | 0.53 | 20.23 | 74.23 | 5869331 | 93.06 | 194197809 | 6494.258402 | 99.21078153 | 236 | 29667 | 549 | 4196 | 13 | 1325 |
| CS0174_1 | 8141485 | 7643857 | 0.43 | 29.33 | 64.04 | 5402144 | 90.61 | 173997072 | 5818.716249 | 99.54854028 | 135 | 29768 | 540 | 2448 | 16 | 876 |
| CS0572_1 | 8997291 | 8171288 | 0.03 | 87.29 | 2.68 | 1038425 | 21.12 | 7790955 | 260.5409156 | 43.20302311 | 16984 | 12919 | 18110 | 21495 | 9 | 18036 |
| CS0175_1 | 10371955 | 9783033 | 0.57 | 1.32 | 97.79 | 9654038 | 99.09 | 339970561 | 11369.11216 | 99.86289001 | 41 | 29862 | 50 | 76 | 13 | 12 |
| CS0635_1 | 13811749 | 13094066 | 0.26 | 59.93 | 35.17 | 5247278 | 87.75 | 163769770 | 5476.700331 | 99.81272782 | 56 | 29847 | 652 | 4223 | 14 | 2056 |
| CS0405_1 | 6992268 | 6600439 | 0 | 96.66 | 0.01 | 220584 | 0.34 | 26824 | 0.8970337424 | 6.75183092 | 27884 | 2019 | 28924 | 29903 | 2 | 28863 |
| CS0406_1 | 9253675 | 8768311 | 0 | 95.22 | 0.06 | 418862 | 1.36 | 202672 | 6.777647728 | 5.912450256 | 28135 | 1768 | 28803 | 29297 | 0 | 28769 |
| CS0636_1 | 9828961 | 9314287 | 0.05 | 86.5 | 8.4 | 1257366 | 62.24 | 27812473 | 930.0897234 | 68.29080694 | 9482 | 20421 | 11650 | 15943 | 9 | 11498 |
| CS0176_1 | 6483938 | 6147561 | 0.4 | 33.12 | 62.56 | 4111493 | 93.55 | 136751625 | 4573.174096 | 99.34120322 | 197 | 29706 | 629 | 4712 | 13 | 1109 |
| CS0637_1 | 7367128 | 6955083 | 0.01 | 93.83 | 1.16 | 428989 | 18.78 | 2864054 | 95.77814935 | 8.142995686 | 27468 | 2435 | 28052 | 28491 | 0 | 25369 |
| CS0177_1 | 8819290 | 8347920 | 0.42 | 30.83 | 60.9 | 5773859 | 88.04 | 180762352 | 6044.957095 | 99.73915661 | 78 | 29825 | 410 | 3611 | 15 | 846 |
| CS0178_1 | 9172347 | 8603805 | 0.39 | 37.98 | 57.12 | 5336363 | 92.1 | 174710764 | 5842.583152 | 99.64886466 | 105 | 29798 | 314 | 2413 | 12 | 1266 |
| CS0638_1 | 8668109 | 8209933 | 0.04 | 87.19 | 5.88 | 1051706 | 45.9 | 17175102 | 574.3604989 | 65.10383574 | 10435 | 19468 | 12882 | 19112 | 20 | 12400 |

| Sample IDs Replicate 2 | In - Depth Custom Pipeline (Replicate 2): Total Reads_Replicate2 | Trimmed Reads q30-30bp_Replicate2 | Hisat2 Human % _Replicate2 | Hisat2 Covidall % _Replicate2 | Hisat2 Unmapped Covid Read#_Replicate2 | Hisat2 Unmapped Covid %_Replicate2 | BasePairescovered_Replicate2 | X coverage _Replicate2 | Genome coverage_Replicate2 | # bases with 0 coverage_Replicate2 | # bases with >0 coverage_Replicate2 | # bases with <10x coverage_Replicate2 | # bases with <100x coverage_Replicate2 | VARSCAN Variant Count_Replicate2 | N's_Replicate2 |
|---|---|---|---|---|---|---|---|---|---|---|---|---|---|---|---|
| CS0169_2 | 8555066 | 7964189 | 25.05 | 72.12 | 5968893 | 96.23 | 204145939 | 6826.938401 | 99.84951343 | 45 | 29858 | 470 | 2269 | 12 | 862 |
| CS0170_2 | 9092791 | 8546744 | 22.26 | 74.97 | 6644229 | 96.43 | 227762653 | 7616.715814 | 99.85285757 | 44 | 29859 | 117 | 640 | 11 | 373 |
| CS0171_2 | 10380360 | 9603719 | 14.68 | 83.38 | 8194277 | 97.72 | 284520148 | 9514.769354 | 99.95987025 | 12 | 29891 | 81 | 399 | 15 | 512 |
| CS0634_2 | 10350054 | 9716680 | 86.01 | 7.15 | 1359482 | 51.1 | 24705434 | 826.1858008 | 79.65755944 | 6083 | 23820 | 9037 | 14936 | 15 | 8551 |
| CS0172_2 | 9004889 | 8494305 | 5.68 | 93.16 | 8011763 | 98.77 | 281330301 | 9408.096211 | 99.93646122 | 19 | 29884 | 77 | 390 | 13 | 337 |
| CS0173_2 | 7711627 | 7221317 | 20.22 | 74.24 | 5761147 | 93.06 | 190577156 | 6373.178477 | 99.17734007 | 246 | 29657 | 510 | 1130 | 13 | 1318 |
| CS0174_2 | 8121883 | 7553322 | 29.29 | 64.07 | 5340997 | 90.6 | 171982727 | 5751.35361 | 99.74250075 | 77 | 29826 | 557 | 1215 | 16 | 878 |
| CS0572_2 | 8764435 | 7819029 | 87.25 | 2.74 | 997120 | 21.51 | 7614626 | 254.6442163 | 43.36019797 | 16937 | 12966 | 18168 | 21509 | 9 | 18080 |
| CS0175_2 | 10323857 | 9638201 | 1.31 | 97.78 | 9512241 | 99.08 | 334839427 | 11197.51955 | 99.85954586 | 42 | 29861 | 50 | 75 | 13 | 12 |
| CS0635_2 | 13690194 | 12832710 | 59.91 | 35.38 | 5145245 | 88.24 | 161448392 | 5399.07006 | 99.84616928 | 46 | 29857 | 257 | 4568 | 15 | 1981 |
| CS0405_2 | 6901212 | 6437374 | 96.6 | 0.01 | 218644 | 0.34 | 26409 | 0.8831555362 | 6.738454336 | 27888 | 2015 | 29062 | 29903 | 1 | 28921 |
| CS0406_2 | 9095576 | 8520606 | 95.13 | 0.07 | 414873 | 1.39 | 205649 | 6.877202956 | 6.052904391 | 28093 | 1810 | 28803 | 29284 | 0 | 28797 |
| CS0636_2 | 9652608 | 9040907 | 85.8 | 8.5 | 1284256 | 59.83 | 27302455 | 913.0339765 | 82.86793967 | 5123 | 24780 | 11658 | 15973 | 9 | 11509 |
| CS0176_2 | 6457441 | 6076960 | 33.01 | 62.6 | 4071086 | 93.45 | 135239048 | 4522.591312 | 99.25759957 | 222 | 29681 | 543 | 3095 | 13 | 1039 |
| CS0637_2 | 7280670 | 6789610 | 93.87 | 1.17 | 416419 | 19.09 | 2825214 | 94.47928302 | 7.661438652 | 27612 | 2291 | 28051 | 28486 | 0 | 28025 |
| CS0177_2 | 8722091 | 8184606 | 30.74 | 61 | 5668261 | 88.08 | 177497173 | 5935.764739 | 99.85620172 | 43 | 29860 | 184 | 478 | 15 | 848 |
| CS0178_2 | 9196970 | 8549922 | 37.96 | 57.17 | 5303953 | 92.15 | 173720039 | 5809.451861 | 99.73915661 | 78 | 29825 | 401 | 2519 | 12 | 1163 |
| CS0638_2 | 8551440 | 8012634 | 86.67 | 5.95 | 1067882 | 44.64 | 16957415 | 567.0807277 | 86.52977962 | 4028 | 25875 | 12975 | 19093 | 18 | 12772 |

| | | RT-PCR | | DRAGEN COVIDSeq Test Pipeline | | In - Depth Custom Pipeline (Merged) | | | | | | | | | | | | | | | In - Depth Custom Pipeline (Replicate 1) | | | | | | | | | | | | | | | | In - Depth Custom Pipeline (Replicate 2) | | | | | | | | | | | | | | |
|---|---|---|---|---|---|---|---|---|---|---|---|---|---|---|---|---|---|---|---|---|---|---|---|---|---|---|---|---|---|---|---|---|---|---|---|---|---|---|---|---|---|---|---|---|---|---|---|---|---|---|---|
| Sample IDs Replicate 1 | Sample IDs Replicate 2 | RT (Pos/Neg) replicate1 | RT (Pos/Neg) replicate 2 | DRAGEN SARS-CoV-2 Detection Replicate 1 | DRAGEN SARS-CoV-2 Detection Replicate 2 | Total Reads_merged | Trimmed Reads q30-30bp_merged | Hisat2 Human % _merged | Hisat2 Covidall % _merged | Hisat2 Unmapped Covid Read#_merged | Hisat2 Unmapped Covid %_merged | BasePairescovered_merged | X coverage _merged | Genome coverage_merged | # bases with 0 coverage_merged | # bases with >0 coverage_merged | # bases with <10x coverage_merged | # bases with <100x coverage_merged | VARSCAN Variant Count_merged | N's_merged | Total Reads_Replicate1 | Trimmed Reads q30-30bp_Replicate1 | Kraken2 coronavirus %_Replicate1 | Hisat2 Human % _Replicate1 | Hisat2 Covidall % _Replicate1 | Hisat2 Unmapped Covid Read#_Replicate1 | Hisat2 Unmapped Covid %_Replicate1 | BasePairescovered_Replicate1 | X coverage _Replicate1 | Genome coverage_Replicate1 | # bases with 0 coverage_Replicate1 | # bases with >0 coverage_Replicate1 | # bases with <10x coverage_Replicate1 | # bases with <100x coverage_Replicate1 | VARSCAN Variant Count_Replicate1 | N's_Replicate1 | Total Reads_Replicate2 | Trimmed Reads q30-30bp_Replicate2 | Hisat2 Human % _Replicate2 | Hisat2 Covidall % _Replicate2 | Hisat2 Unmapped Covid Read#_Replicate2 | Hisat2 Unmapped Covid %_Replicate2 | BasePairescovered_Replicate2 | X coverage _Replicate2 | Genome coverage_Replicate2 | # bases with 0 coverage_Replicate2 | # bases with >0 coverage_Replicate2 | # bases with <10x coverage_Replicate2 | # bases with <100x coverage_Replicate2 | VARSCAN Variant Count_Replicate2 | N's_Replicate2 |
