## Supplementary Table 4a for "High throughput detection and genetic epidemiology of SARS-CoV-2 using COVIDSeq next generation sequencing"

| | COVIDSeq Positive | COVIDSeq Negative | Invalid | TOTAL |
|---|---|---|---|---|
| RT positive | 633 | 16 | 6 | 655 |
| RT Negative | 6 | 13 | 0 | 19 |
| Inconclusive | 21 | 14 | 0 | 35 |
| Pan Sarbeco | 16 | 25 | 2 | 43 |
| **TOTAL** | 676 | 68 | 8 | **752** |

**Supplementary Table 4a**: Summary of the COVIDSeq assay comparison with RT-PCR.
