## Supplementary Table 4b for "High throughput detection and genetic epidemiology of SARS-CoV-2 using COVIDSeq next generation sequencing"

| Isolation Method | RT-PCR Postive | | | RT-PCR Negative | | | pan-sarbeco | | | Inconclusive | | | Total |
|---|---|---|---|---|---|---|---|---|---|---|---|---|---|
| | COVIDSeq Detected | COVIDSeq Not Detected | COVIDSeq Invalid | COVIDSeq Detected | COVIDSeq Not Detected | COVIDSeq Invalid | COVIDSeq Detected | COVIDSeq Not Detected | COVIDSeq Invalid | COVIDSeq Detected | COVIDSeq Not Detected | COVIDSeq Invalid | |
| Nextractor | 263 | 1 | 0 | 4 | 7 | 0 | 12 | 19 | 0 | 13 | 3 | 0 | 322 |
| Trueprep | 195 | 5 | 1 | 1 | 5 | 0 | 3 | 4 | 2 | 5 | 7 | 0 | 228 |
| QIAamp | 168 | 10 | 4 | 1 | 1 | 0 | 1 | 2 | 0 | 3 | 4 | 0 | 194 |
| TRizol | 7 | 0 | 1 | 0 | 0 | 0 | 0 | 0 | 0 | 0 | 0 | 0 | 8 |
| Total | **633** | **16** | **6** | **6** | **13** | **0** | **16** | **25** | **2** | **21** | **14** | **0** | **752** |

**Supplementary Table 4b:** Comparison of different RNA extraction methods and detection of the SARS-CoV-2 with RT-PCR and COVIDSeq test.
